# Stage-Specific Threats Reveal the Inadequacy of Adult-Centered Conservation

**DOI:** 10.64898/2026.03.17.712346

**Authors:** Yanfang Song, Yongle Wang, Qingqing Li, Zhiyong Yuan, Weiwei Zhou

## Abstract

In an era of severe global biodiversity threats, understanding the link between species’ traits and their endangerment helps uncover causes of risk and infer threats to understudied species. Most animals have complex life cycles with distinct stages that may face stage-specific threats. Current conservation frameworks rely heavily on adult traits, potentially misjudging extinction risk. Using Chinese anurans as a model, we integrated functional traits from both adult and tadpole stages to examine their association with extinction risk. We found that body size positively correlates with risk in both stages. Microhabitat use related with extinction risk in tadpoles but shows no significant link in adults. Adult relative tympanum diameter and head length also correlate with extinction risk. These results indicate that species vulnerability is correlated with multi-stage traits, with both shared and stage-specific threats. Conservation based solely on adult traits may fail to accurately assess species threats. We call for integrating a whole-life-history perspective into biodiversity assessment and conservation to more effectively address the global biodiversity crisis.

## 1 Introduction

Human activities are driving global biodiversity loss at an unprecedented rate and scale, placing a vast number of species at risk of extinction (Cowie et al., 2022; Keck et al., 2025). However, not all species are affected equally or in the same manner. Species with different ecological strategies exhibit marked differences in vulnerability when facing the anthropogenic disturbances (Chichorro et al., 2022). Although the statistical associations between traits and extinction risk may not imply causation, investigating the relationship between species’ traits and extinction risk is of great importance for revealing causes of endangerment and predicting potential threats. On one hand, understanding the relationship between traits and extinction risk helps identify species’ characteristics that may make it vulnerable, thereby clarifying the causes of their endangerment. For example, body size, as the most fundamental and commonly measured ecological trait, has been shown to be closely related to extinction risk across numerous taxa (Tomiya, 2013; Vilela et al., 2014; Nolte et al., 2019; Cardillo, 2021). The relationships between body sizes and extinction risks vary among different groups. Large species often face higher risks due to low reproductive rates, narrow distributions, or greater susceptibility to human hunting (Keane et al., 2005; Ruland & Jeschke, 2017). But in some taxa, small species may be more prone to extinction due to smaller geographic ranges, weak dispersal ability or habitat specialization (Owens & Bennett, 2000; Cardillo, 2021). On the other hand, our knowledge of biodiversity remains incomplete. Even for vertebrate groups, a large number of new species are described annually (Uetz et al., 2025; Frost & Darrel, 2026). Existing assessment systems struggle to promptly evaluate the extinction risk of these new species. Understanding the relationship between traits and extinction risk can provide a basis for risk prediction for species that are not yet assessed or are data-deficient (González-del-Pliego et al., 2019).

Exploring the relationship between species’ functional traits and extinction risk is very important for biodiversity conservation (Chichorro et al., 2019). However, a long-overlooked issue is that over 80% of described animal species have complex life cycles (Werner, 1988; Phung et al., 2020). In these groups, different life stages are often highly differentiated in morphology, physiology, behavior, and habitat requirements (Moran & Nancy, 1994). According to the adaptive decoupling hypothesis, trait evolution in these stages often faces different selective pressors and shows relative independence (Wollenberg Valero et al., 2017). This hypothesis has been supported in many groups (Phillips, 1998; Sherratt et al., 2017; Goedert & Calsbeek, 2019). It follows that the threats and vulnerabilities a species faces at different stages may also differ. Unfortunately, studies on drivers of extinction risk always rely on adult traits, with minimal attention paid to juvenile or non-adult stages, leading to potential systematic bias in our understanding of a species’ overall vulnerability. For instance, in amphibians, which is a typical group with biphasic life cycles, conservation measures that neglect larval stages are often less effective (Nolan et al., 2023). Furthermore, studies have shown that biodiversity hotspots for tadpoles and adults do not overlap, and conservation priority species defined based on a single stage may have great biases in amphibians (Song et al., 2025a). Meanwhile, incorporating traits from life stages other than adults into models allows for the development of more robust statistical associations between traits and threat status, thereby enhancing the effectiveness of biodiversity conservation efforts.

Anuran amphibians serve as an ideal model system for investigating how functional traits across different life stages relate to extinction risk (Phung et al., 2020). As a group characterized by complex life cycles, amphibians exhibit striking differences between adults and their larval stage, also known as tadpoles, in morphology, diet, microhabitat selection, and behavior (McDiarmid & Altig, 1999). Furthermore, with over 40% of amphibian species facing extinction risk, they rank among the most threatened vertebrate groups (IUCN, 2025). While population models suggest that survival during the tadpole stage may not be the most critical determinant of overall population persistence (Biek et al., 2002), and other studies have highlighted the importance of the post-metamorphic juvenile stage (Petrovan & Schmidt, 2009), the tadpole stage nevertheless remains a vulnerable phase in amphibian development, characterized by high mortality and sensitivity to disturbances (Nolan et al., 2023). Consequently, insufficient understanding of this life stage may hinder effective conservation of the entire group (Nolan et al., 2023). Examining the relationship between functional traits at different life stages and extinction risk in amphibians not only aids in protecting this unique and endangered group but also offers valuable insights for other taxa with complex life histories. Regrettably, most existing research has focused primarily on adult traits (Cooper et al., 2008; Chen et al., 2019; Cardillo, 2021). Although some studies incorporate reproductive traits, the link between tadpole-stage traits and extinction risk remains poorly understood (Chen et al., 2019). At the same time, although tadpole conservation has received attention in some amphibian conservation efforts (Calhoun et al., 2014; Moor et al., 2022; Vredenburg, 2004), current conservation assessment frameworks and conservation planning still generally lack the integration of information on this life stage, which may reduce the effectiveness of these conservation measures. This limitation observed in amphibians may be widespread across most animal groups with complex life cycles (Faria et al., 2021). Therefore, using amphibians as a model, the research framework considering information from different life history stages holds broad implications for advancing the conservation of complex life cycles taxa throughout the animal kingdom. It helps address the bias in current conservation programs which are heavily relying on information of adults.

For groups with complex life histories, the lack of data on stages other than adults severely limits the related research. China harbors exceptional amphibian diversity and has accumulated comprehensive, systematic data on species taxonomy, distribution, ecology, and conservation status (Fei et al., 2009a; Fei et al., 2009b; Fei, 2020; Huang et al., 2023; Song et al., 2025a; Song et al., 2025b; Wei et al., 2025; AmphibiaChina, 2026), providing a unique opportunity for integrated analyses of multi-stage traits and extinction risk. Critically, these data include high-resolution continuous functional trait data, which, compared to traditional discrete or categorical variables, can more finely depict variation in species’ ecological strategies (Tobias et al., 2022). Based on high-quality data, we will use Chinese anuran species to address two questions by studying the relationship between extinction risk and functional traits of adults and larvae: (1) Are functional traits of both adults and tadpoles correlated with species extinction risk? (2) If correlated, are these associations consistent across different life stages? By comparing correlations between traits and extinction risk across different stages, we aim to evaluate whether the adult-based conservation assessment system adequately reflects the vulnerability of species with complex life histories.

## 2 Methods

### 2.1 Data Collection

Extinction risk data were sourced from the China Biodiversity Red List: Vertebrates Volume (2020) (MEP & CAS, 2023), which is the latest species threat category list for species in China currently. Compared to the International Union for Conservation of Nature (IUCN) Red List, this list is updated more timely for Chinese species and additionally covers the threat categories of 42 species. Similar to IUCN Red List, the China Biodiversity Red List categories are assigned based on population size and distribution range, and are independent of species’ traits. We also assessed the similarity of species assessments between the two lists based on Spearman correlation and reanalyzed the data based on IUCN Red List. Since our aim was to examine relationships between traits and threat status, rather than predicting the threat status of unevaluated species, we excluded species that were data deficient. Following the method of previous studies (Cardillo, 2021), we performed an ordered coding conversion according to the severity of species threat categories: Least Concern (LC) = 0; Near Threatened (NT) = 1; Vulnerable (VU) = 2; Endangered (EN) = 3; Critically Endangered (CR) = 4.

Functional traits can reflect species’ interactions with the environment (Violle et al., 2007). We selected eight morphological traits of both adults and tadpole related to locomotion, foraging, and ecological habits from functional trait databases (Huang et al., 2023; Song et al., 2025b). These traits are closely related to the ecological function and performance of species (Table 1). Tadpole traits were collected specifically during Gosner stages 32–40, when taxonomically diagnostic characters remain stable (Haas et al., 2022). Except for body size, all other morphological data were converted to ratios relative to total length of tadpoles or snout-vent length of adults. In addition to morphological traits, previous studies have shown that microhabitat type is a strong predictor of extinction risk (Seibold et al., 2015). Therefore, we also included microhabitat data for both adults and tadpoles, encoded as dummy variables. Although amphibians typically breed in aquatic environments, most species spend most time of their lives outside water. Thus, microhabitat here refers to the primary non-breeding habitat. Adult microhabitats were classified as: arboreal, fossorial, terrestrial, aquatic, and torrential. Tadpole microhabitats were categorized as lentic or lotic. If a species can inhabit multiple microhabitats, all applicable types were annotated. For example, semi-aquatic was coded as being able to inhabit both terrestrial and aquatic habitats. As our purpose was to compare the relationships between adults’ and tadpoles’ traits and the extinction risk, we did not include traits that have been shown to be closely associated with extinction risk, such as geographic range, elevational range, fecundity, and habitat specificity.

**Table 1:**
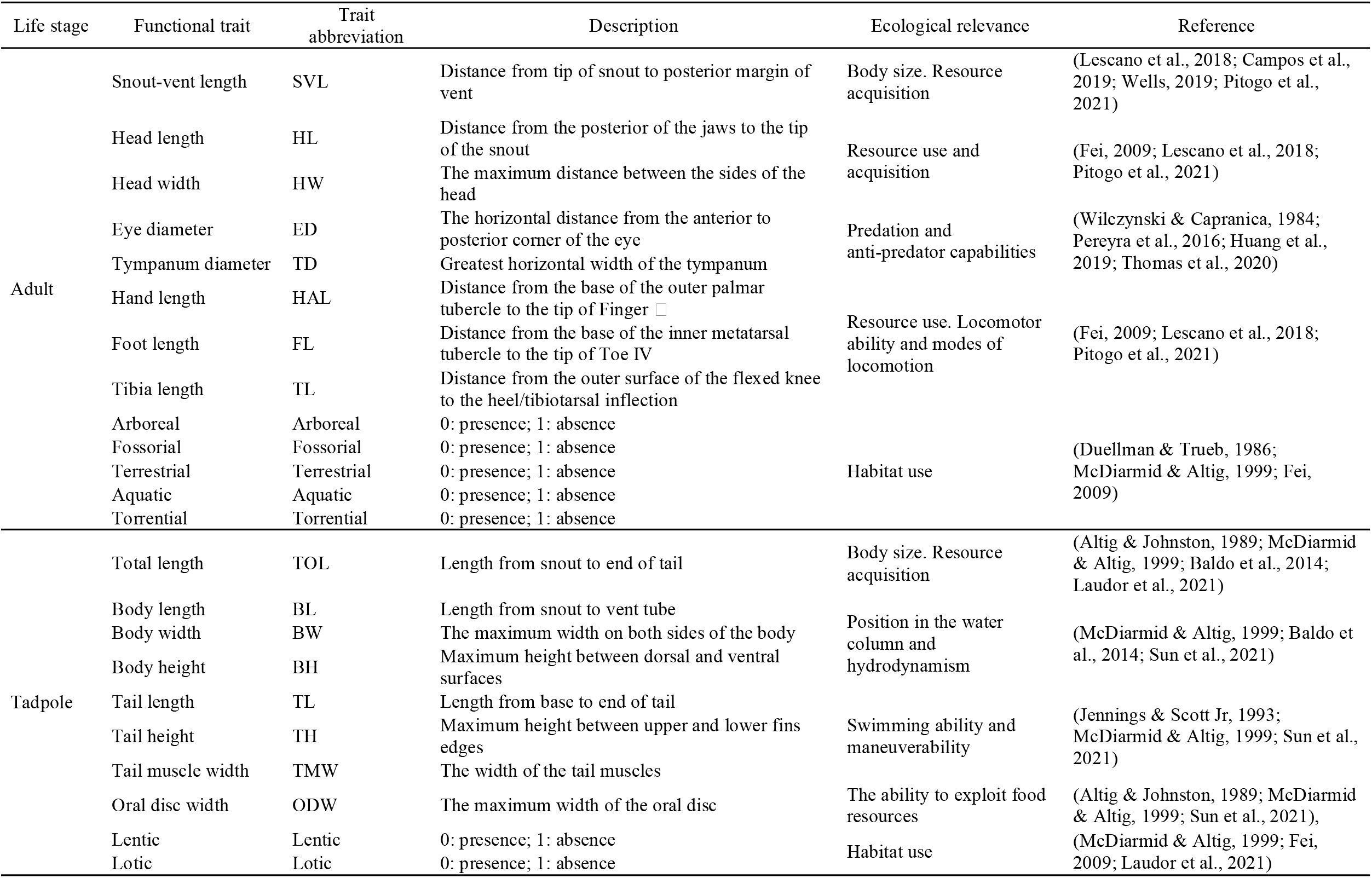
Functional traits and their potential ecological functions.

Our compiled dataset includes functional trait data for 375 species. The completeness of the trait data is very high, morphological traits ranging from 68.7% to 100%, and information of microhabitat covered all adults and 76.5% of tadpoles (Fig. S1). We performed data imputation considering phylogenetic relationships. The phylogenetic data we used came from previous studies (Xu et al., 2024; Song et al., 2025a), covering all Chinese anuran species. First, we calculated the phylogenetic distance matrix for the species based on the phylogenetic tree. Then, through principal coordinates analysis, we extracted the principal coordinates of the phylogenetic distance matrix, merged them with the morphological traits and microhabitat data. In total, we used 100 axes, which could account 99.4 % of the variances. Missing values were imputed using the R package missForest (Stekhoven, 2022), a random forest–based method proven highly accurate for high–dimensional data with complex, nonlinear interactions (Stekhoven & Bühlmann, 2011; Penone et al., 2014). We used its default configuration: maximum iterations were set to 10; initial fill values used the median for numerical variables and the mode for categorical variables; the random forest contained 100 decision trees, with a minimum leaf node sample size of 1, a minimum sample size for internal node splitting of 2. The algorithm iteratively refines predictions across variables until convergence or the maximum iteration limit is reached.

### 2.2 Data Analysis

Prior to analysis, we assessed multicollinearity among all trait variables using variance inflation factors (VIFs). The VIF for each variable was below 5, indicating acceptable levels of collinearity (Table S1). We employed phylogenetic generalized least squares (PGLS) models to examine associations between species traits and threat categories. To account for non-independence due to shared evolutionary history, we incorporated a phylogenetic covariance matrix derived from the species tree. Following the method of Swenson (2014) and Revell et al (2022), models were fitted in R using the gls function from the *nlme* package (Pinheiro et al., 2025), under both Brownian motion (BM) and Ornstein–Uhlenbeck (OU) models of trait evolution. To assess the impact of phylogenetic structure on model performance, we also fitted non-phylogenetic generalized least squares (GLS) models. Model performance was compared using the small-sample corrected Akaike Information Criterion (AICc), calculated with the MuMIn package (Bartoń, 2023).

To further ensure the robustness of our model results, we also conducted a multi-model inference analysis using the dredge function in the MuMIn package (Bartoń, 2023) based on the best-fitting evolutionary model, and calculated trait importance. By comparing variable importance rankings and the full models between the adult and larval stages, we further assessed how traits from different life stages differ in predicting extinction risk. We also repeated the analysis based on the categories of the IUCN Red List.

Additionally, to test if body size of adults and tadpoles were correlated, we examined the correlation between adult snout-vent length and tadpole total length. To correct for phylogenetic non-independence in correlation estimates, we used the corphylo function from the ape package to compute phylogenetically informed Pearson correlations (Paradis & Schliep, 2018). We first assessed this relationship across all species, then repeated the analysis within families containing more than five species.

All data compilation, analysis, and visualization were performed in R version 4.4.1 (R Core Team, 2024). Original files and code are accessible at figshare (https://doi.org/10.6084/m9.figshare.31149463).

## 3 Results

### 3.1 Threat Status of Chinese Anurans

After excluding species not evaluated or data deficient in the China Biodiversity Red List, 299 species were included in the analysis. Threat status varied markedly among families (Fig. 1). Dicroglossidae exhibited the highest level of threat: 58% of species were classified as Vulnerable (VU) and 18% as Endangered (EN). Megophryidae also faced high threats, with nearly half (48%) of species in threatened categories. A total of 2% of species were classified as Critically Endangered, 30% as VU, and 16% as EN. In Bufonidae, Ranidae, and Rhacophoridae, the proportion of threatened species were 22%, 25% and 24%, respectively. In contrast, Hylidae and Microhylidae consisted almost entirely of Least Concern (LC) species, with no threatened taxa recorded. Bombinatoridae had no CR or EN species, but 33% species were listed as VU. The results of the Spearman test support a significant correlation between the threat categories assigned in the China Biodiversity Red List and those in the IUCN Red List (Spearman’s ρ=0.573, *p* < 0.001, n = 257).

**Figure 1.**
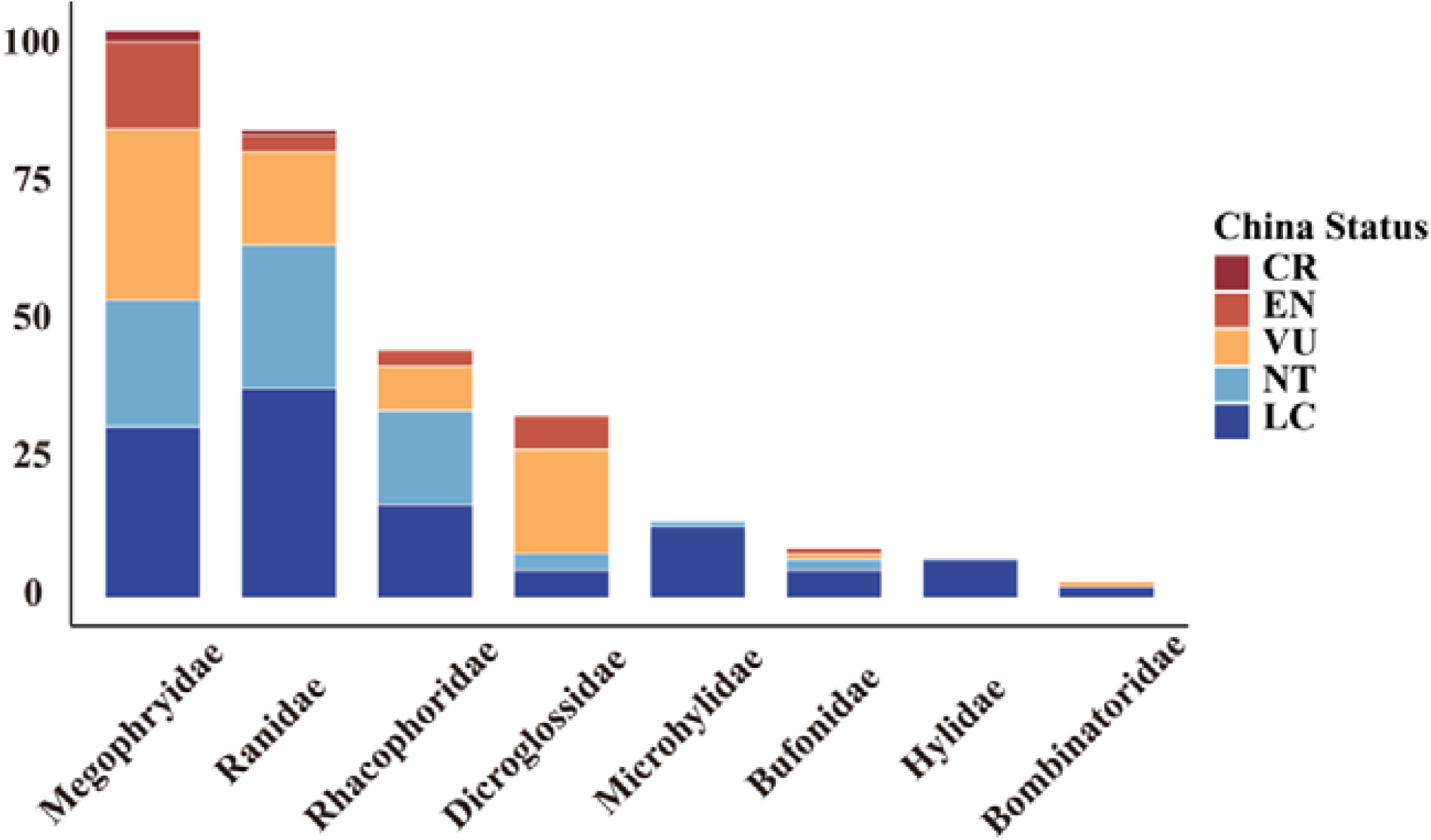
Distribution of threat status across anuran families in China, based on the China Biodiversity Red List– Vertebrates Volume (2020). Each stacked bar represents the proportion of species within a family categorized as Critically Endangered (CR), Endangered (EN), Vulnerable (VU), Near Threatened (NT), or Least Concern (LC). Families are ordered by total number of species.

### 3.2 Extinction Risk Correlates in Adults and Tadpoles

Model comparisons revealed that the OU-based model provided the best fit for both adult and tadpole datasets (Table 2). In adults, snout-vent length, head length, and tympanum diameter showed significant associations with threat status (Fig. 2A). Specifically, snout-vent length (β = 0.52, p < 0.01) and head length (β = 6.86, p < 0.001) were positively correlated with threat level, whereas tympanum diameter (β = −5.91, p < 0.05) was negatively correlated. Other morphological variables such as head width, eye diameter, and traits about forelimb and hindlimb showed no significant effects. Notably, adult microhabitat types were not significantly associated with extinction risk. Furthermore, the analysis based on model averaging yielded similar results (Fig. S2). HL, SVL and TD were the three most important variables to explain extinction risk.

**Table 2.**
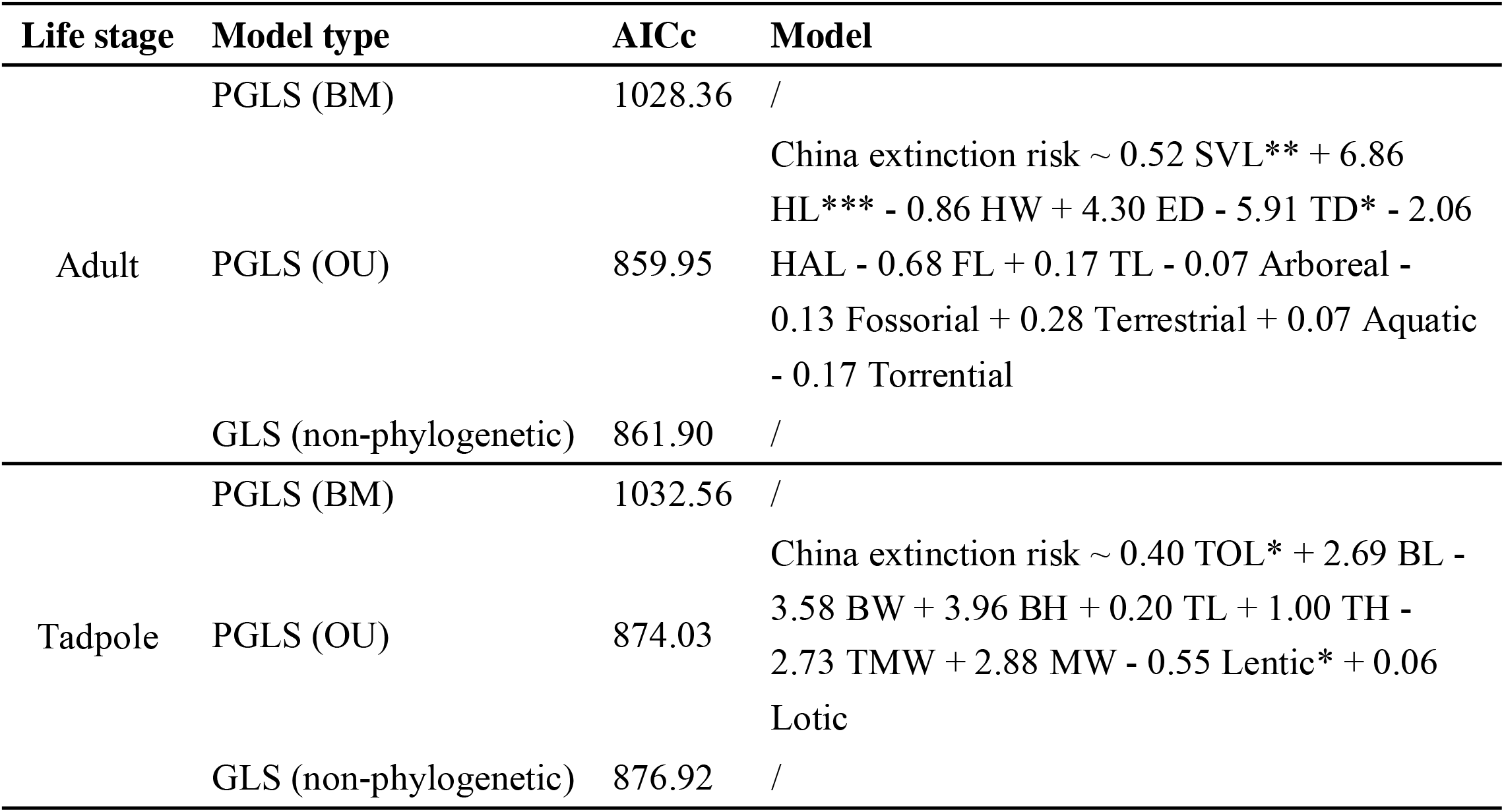
Regression analysis results of extinction risk for adult and tadpole stages. For abbreviations of traits, please refer to Table 1. BM, Brownian motion; OU, Ornstein-Uhlenbeck; GLS, generalized least squares; PGLS, phylogenetic generalized least squares; AICc, Akaike Information Criterion corrected for small sample size.

**Figure 2.**
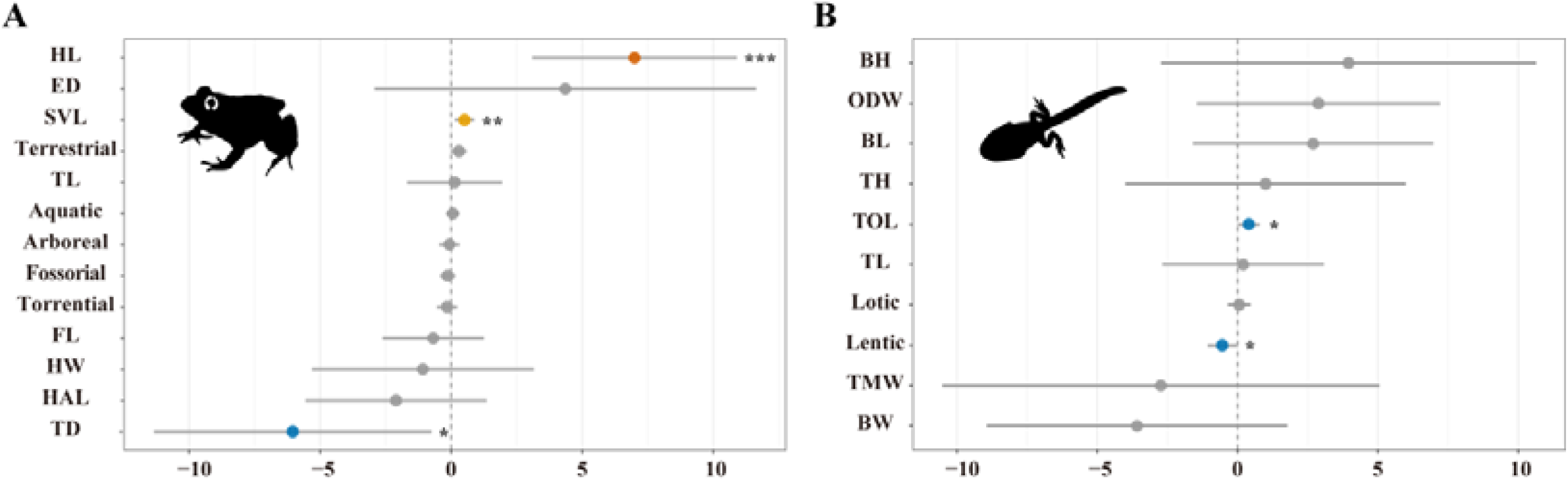
Phylogenetically corrected associations between functional traits and extinction risk in Chinese anurans, separately for (A) adults and (B) tadpoles. Each dot represents the PGLS regression coefficient (β) for a given trait, with horizontal lines indicating 95% confidence intervals. Traits significantly associated with threat status are marked with asterisks (*p < 0.05; **p < 0.01; ***p < 0.001). For abbreviations of traits, please refer to Table 1. The silhouette images are sourced from https://www.phylopic.org.

In tadpoles, only total length and lentic microhabitat use showed significant associations (Fig. 2B). Similar to adults, tadpole body size had a significant positive effect (β = 0.40, p < 0.05). Lentic habitat type showed a significant negative correlation (β = -0.55, p < 0.05), indicating that species which have tadpoles living in lentic environments had relatively lower extinction risk. None of the other traits showed significant correlations. In the model averaging analysis, the two variables representing habitat were the most important, and the third most important was total length, which represents body size. Correlation between adult and tadpole body size was very weak across all species (r = 0.12), with variation among families (Table 3). In the analysis based on IUCN Red List data, the regression coefficients for each trait were similar, but some traits’ coefficients were not significant, including adult head length, body size, and tadpole microhabitat type (Fig. S3 & Fig. S4, Table S2).

**Table 3.** Relationships between tadpole and adult size within anuran families. SVL: snout-vent length; TOL: total length. d: parameter quantifying the rate of return to the optimal value in the Ornstein-Uhlenbeck (OU) evolutionary model, reflecting the strength of phylogenetic signal.

| **Family** | **Coefficient** | **d (SVL)** | **d (TOL)** |
| --- | --- | --- | --- |
| All | 0.12 | 0.58 | 0.13 |
| Bufonidae | 0.11 | 0.00 | 2.74 |
| Dicroglossidae | 0.30 | 0.79 | 0.06 |
| Hylidae | -0.12 | 0.22 | 1.47 |
| Megophryidae | 0.33 | 0.67 | 0.09 |
| Microhylidae | 0.37 | 1.93 | 0.53 |
| Ranidae | 0.34 | 0.00 | 0.00 |
| Rhacophoridae | 0.24 | 0.50 | 0.55 |

## 4 Discussion

By integrating functional trait data from both adult and tadpole stages, we reveal statistical associations between multi-stage traits and extinction risk in anurans. The results support that species threat status is related with traits from both life history stages, and the nature of these associations can be different between stages. Given that over 80% of described animal species have complex life histories (Werner, 1988; Phung et al., 2020), and current conservation programs heavily rely on adult information (Faria et al., 2021), our study suggests that neglecting the vulnerability of non-adult stages may lead to a systematic underestimation of a species’ overall extinction risk. Therefore, to better protect biodiversity, multi-stage perspectives must be incorporated into assessment frameworks and conservation planning (Song et al., 2025a; Roehrdanz, 2026).

### 4.1 Differences in Threats Across Life Stages

Our results confirm that different life stages face distinct threats. Our analysis clearly showed that tadpole microhabitat type was significantly correlated with extinction risk, while adult microhabitat type showed no such association. Although the results of the model averaging analysis slightly differed from those of the full model, both supported that tadpole microhabitat type is associated with extinction risk. Lotic microhabitat was also an important variable explaining extinction risk in the model averaging analysis. Although it was not significant in the full model, the regression coefficient between lotic microhabitat and extinction risk was positive. All the evidences support species with tadpoles living in lentic environments (e.g., ponds) faced significantly lower extinction risk. Habitat alteration caused by human activities is a major threat to biodiversity (Keck et al., 2025). Our results indicate that the impact of habitat change differs across life stages. Therefore, focusing only on one stage, such as adults, may underestimate the threats to biodiversity. Tadpoles are more dependent on aquatic environments than adults (McDiarmid & Altig, 1999), making them particularly sensitive to waterbody degradation (Nolan et al., 2023). The significantly lower extinction threat for lentic tadpoles may be because human-induced habitat changes have a lesser impact on lentic habitats. For example, human-dominated environments, including cities and farmlands, are more likely to retain lentic water bodies, such as ponds in urban parks or farmland (Hamer & Parris, 2011; Holzer, 2014). This result suggests that in amphibian conservation, more attention should be paid to protecting larval habitats. We also recommend that tadpole microhabitat sensitivity be considered as an auxiliary indicator in future species extinction risk assessments.

Apart from body size and microhabitats, no other functional traits in tadpoles showed a significant relationship with extinction risk. However, for adults, besides body size, relative tympanum size and head length were also significantly correlated with extinction risk. We found that species with relatively smaller tympanum faced higher extinction risk. Tympanum size is related to acoustic communication in amphibians, and most anurans rely heavily on acoustic communication for reproduction (Schwartz et al., 2001). Studies have shown that anthropogenic noise causes auditory masking in frogs and reduces individual reproductive success (Caorsi et al., 2019; Grenat et al., 2019; Grenat et al., 2024). It also alters community-level acoustic characteristics (Zhao et al., 2025a). However, this evidence is largely limited to behavioral changes, and no study has directly linked noise to amphibian population viability. Additionally, the IUCN does not recognize noise as a major threat to amphibians. Thus, the link between tympanum size and extinction risk is proposed as a testable hypothesis, and the causal mechanisms need future verification.

We also detected a significant positive correlation between relative head length and extinction risk. Species with larger relative head length had greater extinction risk. Skull morphology is related to both habitat selection and diets in anurans (Paluh et al., 2020). For example, fossorial species tend to have short, high skulls, while aquatic species often possess elongated, flattened heads. Unfortunately, for the species involved in our study, research on the threat differences faced by species with different relative head lengths is still lacking. Simultaneously, we found that relative head length varied greatly among different families (Fig. S5). In several families with high species richness and many endangered species, relative head length was larger, such as in Megophryidae and Dicroglossidae. Therefore, we cannot completely rule out the possibility that the relationship between relative head length and extinction risk is because species in these families have larger relative head lengths. Although we corrected for the influence of phylogenetic relationships in the analysis, strong coupling between taxonomic group characteristics and threat status may still leave a signal. Although the underlying mechanism remains unclear, which warrants further investigation, our study discovered a potential link between adult head length and extinction risk.

Our results clearly indicate that threats differ between adult and tadpole stages in amphibians. These differences suggest that characteristics of different life stages should receive attention in biodiversity conservation. For example, the impact of various human activities on tadpole habitats should receive more focus. Furthermore, it is highly likely that such differences exist in other groups with complex life cycles. Although the vast majority of described animal species have complex life histories, we have a widespread knowledge gap regarding all aspects of stages other than adults (Faria et al., 2021; Nori et al., 2025). This gap may greatly hinder biodiversity conservation policy-making and related research.

### 4.2 Common Threats Across Life Stages

Despite stage differences in threats, we also identified a common association throughout the life cycle: larger body size is associated with higher extinction risk. Significant positive correlations were observed in both adults and tadpoles. Among various functional traits, body size is the most commonly associated functional traits with extinction risks (Tomiya, 2013; Seibold et al., 2015; Terzopoulou et al., 2015; Verde Arregoitia, 2016). Several mechanisms may explain this pattern. Large species are often preferentially targeted by humans for consumption or trade (Terzopoulou et al., 2015; Verde Arregoitia, 2016; Chichorro et al., 2019). In China, large frogs such as *Quasipaa spinosa, Hoplobatrachus chinensis*, and *Rana dybowskii* have suffered severe population declines due to overharvesting (Fei et al., 2009b; Tian et al., 2011; Chan et al., 2014; Zhao et al., 2014). Our results suggest that human harvesting may also threaten species at tadpole stage. Although less common than adults, in many regions tadpoles are also collected for food or traditional medicine (Luo et al., 2023; Liu et al., 2024). Furthermore, smaller-bodied species typically exhibit higher natural abundance (Damuth, 1981), which may provide them with a certain buffer against extinction risk. Alternatively, as body size is related to a species’ resource acquisition, larger species may require more space or resources (Purvis et al., 2000; Chichorro et al., 2019). When the environment is changed or habitat becomes fragmented, difficulties in resource accessibility may also cause species endangerment (Womersley et al., 2024; Ning et al., 2025; Zhao et al., 2025b). This could also be a reason for the positive correlation between body size and extinction risk. Regardless of the exact mechanism, the consistent positive association between size and risk has practical utility. In the absence of detailed life history data, body size can serve as a proxy variable for preliminary risk screening for Chinese amphibians. In extinction risk assessments or conservation priority ranking, species with larger body size, whether adults or tadpoles, should receive more attention.

Of course, we also cannot exclude another hypothesis, which is that adult and tadpole body sizes are correlated. Previous study found a weak correlation between adult and tadpole body sizes, but this correlation varies among different groups (Phung et al., 2020). Among the species involved in this study, adult and larval body sizes showed only a very weak correlation, with a correlation coefficient of only 0.12 (Table 3). Therefore, although we cannot completely rule out the possibility that the relationship between tadpole body size and extinction risk is due to a correlation between tadpole and adult body sizes, our study supports a conservative inference: species possessing larger tadpoles overall face higher risk, and this risk should not be neglected in conservation planning.

## Conclusion

This study demonstrates that extinction risk in Chinese anurans is shaped by functional traits across multiple life stages. On one hand, stage-specific traits reveal unique combinations of threats faced at different life stages. On the other hand, body size, as a cross-stage common trait, is associated with higher extinction risk in both stages. These findings emphasize that focusing solely on the adult stage will underestimate the overall extinction risks of species, limit our understanding of the threats species face, and reduce the effectiveness of biodiversity conservation measures. Based on our results, we recommend incorporating information from different life history stages when assessing species threat status. Although this study is based on anuran species, stage-specific vulnerability likely has broad applicability. Given the large amounts of animal species have complex life histories, we suggest integrating information across all life stages to build more effective frameworks for global biodiversity conservation.

## Acknowledgements

This work was supported by the funding agencies, for which we are deeply grateful. This work was funded by the National Natural Science Foundation of China (NSFC32170445) and Science and Technology Projects of Xizang Autonomous Region, China (XZ202402ZD0005) to Weiwei Zhou. National Natural Science Foundation of China (NSFC31860624) to Qingqing Li. National Natural Science Foundation of China (NSFC32570496; NSFC32370478) to Zhiyong Yuan.

## Supplementary Information

**Table S1.** The results of variance inflation factors for adult and tadpole traits. For abbreviations of traits, please refer to Table 1.

| Life stage | Trait | VIF value |
| --- | --- | --- |
| Adult | SVL | 1.5 |
|  | HL | 2.0 |
|  | HW | 2.1 |
|  | ED | 1.5 |
|  | TD | 1.4 |
|  | HAL | 1.1 |
|  | FL | 1.5 |
|  | TL | 1.6 |
| Tadpole | TOL | 1.2 |
|  | SVL | 2.9 |
|  | BW | 3.6 |
|  | BH | 4.5 |
|  | TL | 1.9 |
|  | TH | 2.2 |
|  | TMW | 1.3 |
|  | MW | 1.3 |

**Table S2.**
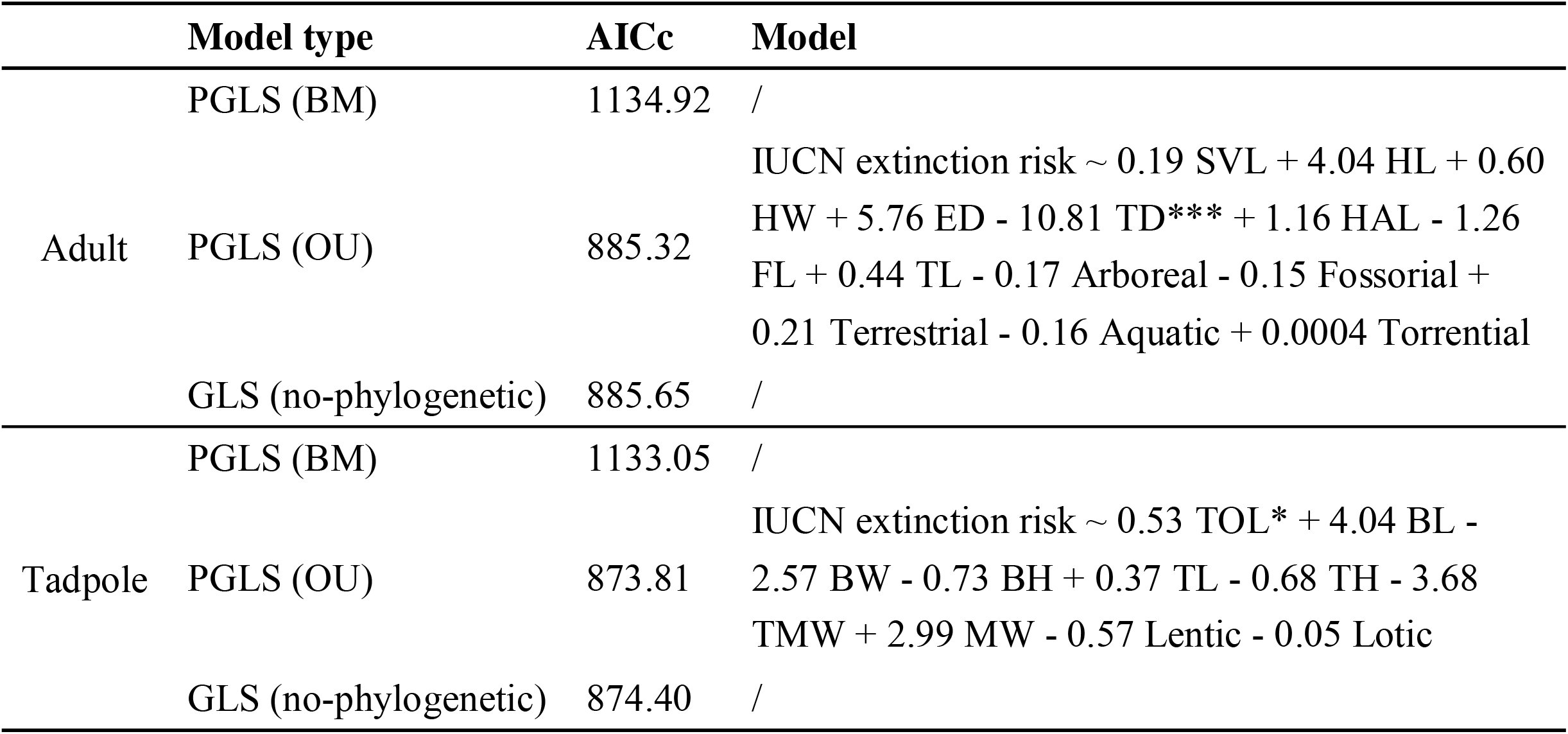
Regression analysis results of extinction risk for adult and tadpole stages (IUCN). For abbreviations of traits, please refer to Table 1. BM, Brownian motion; OU, Ornstein–Uhlenbeck; GLS, generalized least squares; PGLS, phylogenetic generalized least squares; AICc, Akaike Information Criterion corrected for small sample size.

**Figure S1.**
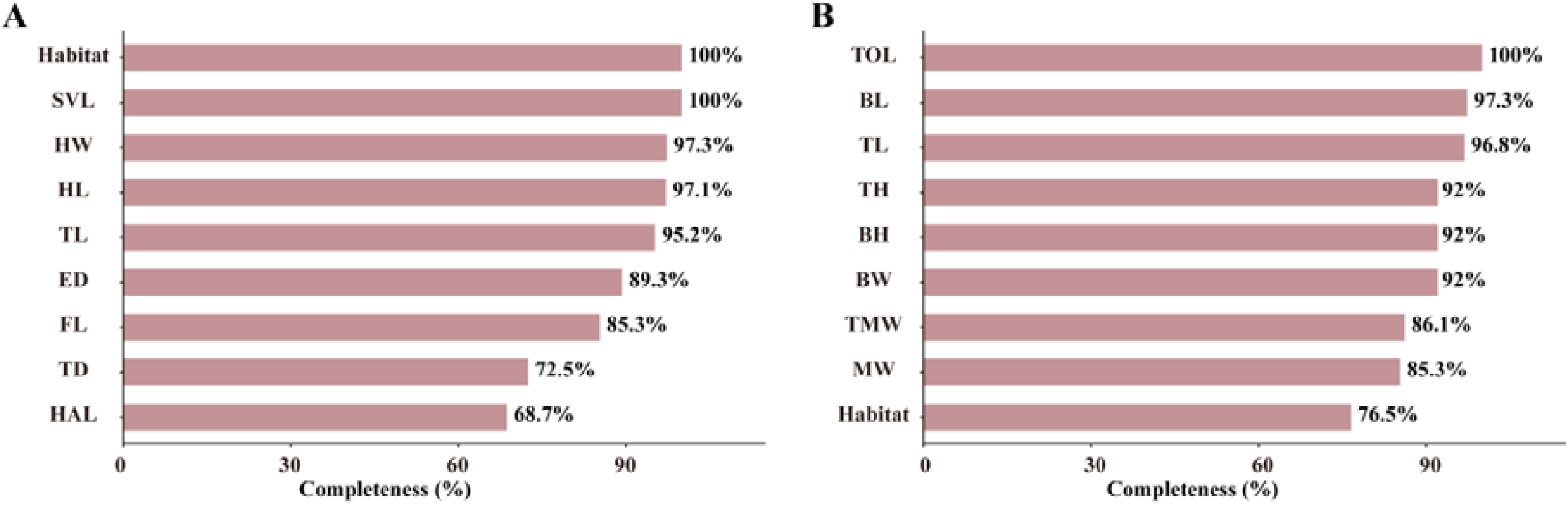
Data completeness of each trait for (A) adults and (B) tadpoles.

**Figure S2.**
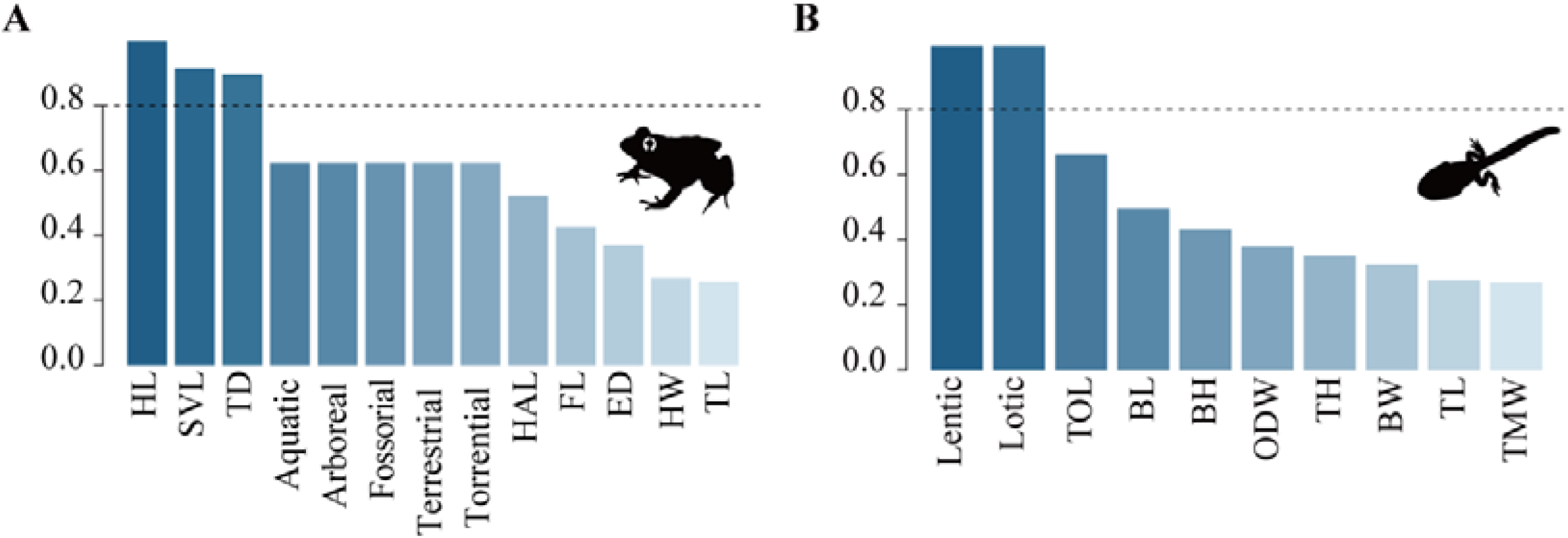
Trait importance (Sum of Akaike Weights) of adult and tadpole traits based on model averaging under the OU evolutionary model using China Biodiversity Red List. For abbreviations of traits, please refer to Table 1. The silhouette images are sourced from https://www.phylopic.org.

**Figure S3.**
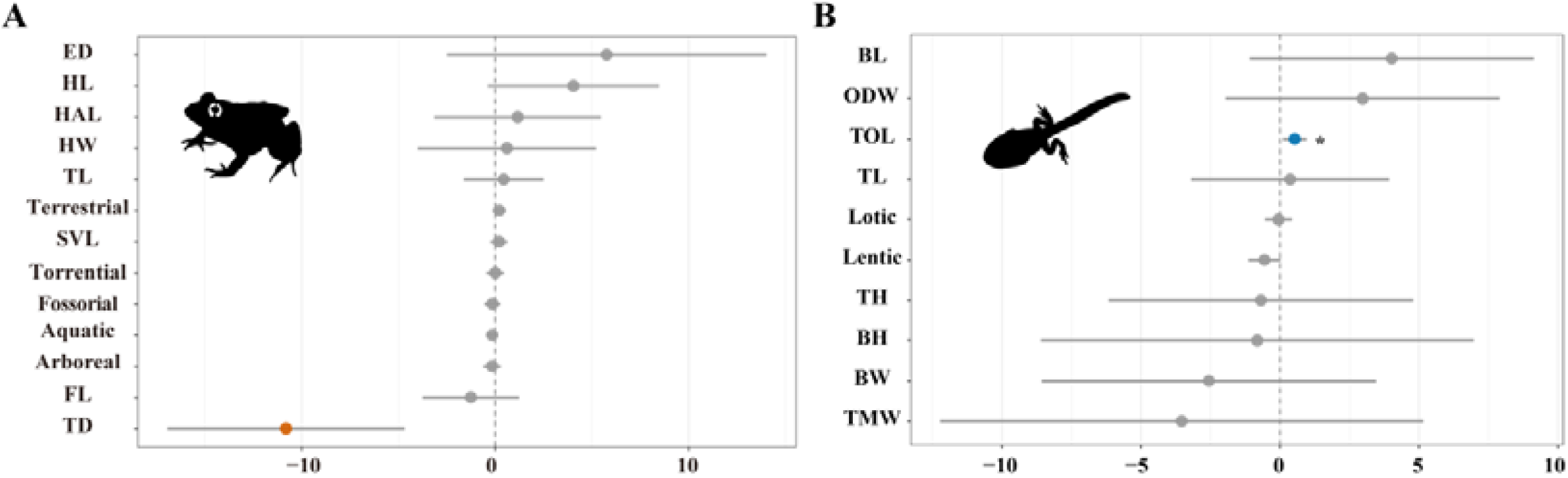
Phylogenetically corrected associations between functional traits and extinction risk in Chinese anurans based on IUCN Red list for (A) adults and (B) tadpoles. Each dot represents the PGLS regression coefficient (β) for a given trait, with horizontal lines indicating 95% confidence intervals. Traits significantly associated with threat status are marked with asterisks (*p < 0.05; **p < 0.01; ***p < 0.001). For abbreviations of traits, please refer to Table 1. The silhouette images are sourced from https://www.phylopic.org.

**Figure S4.**
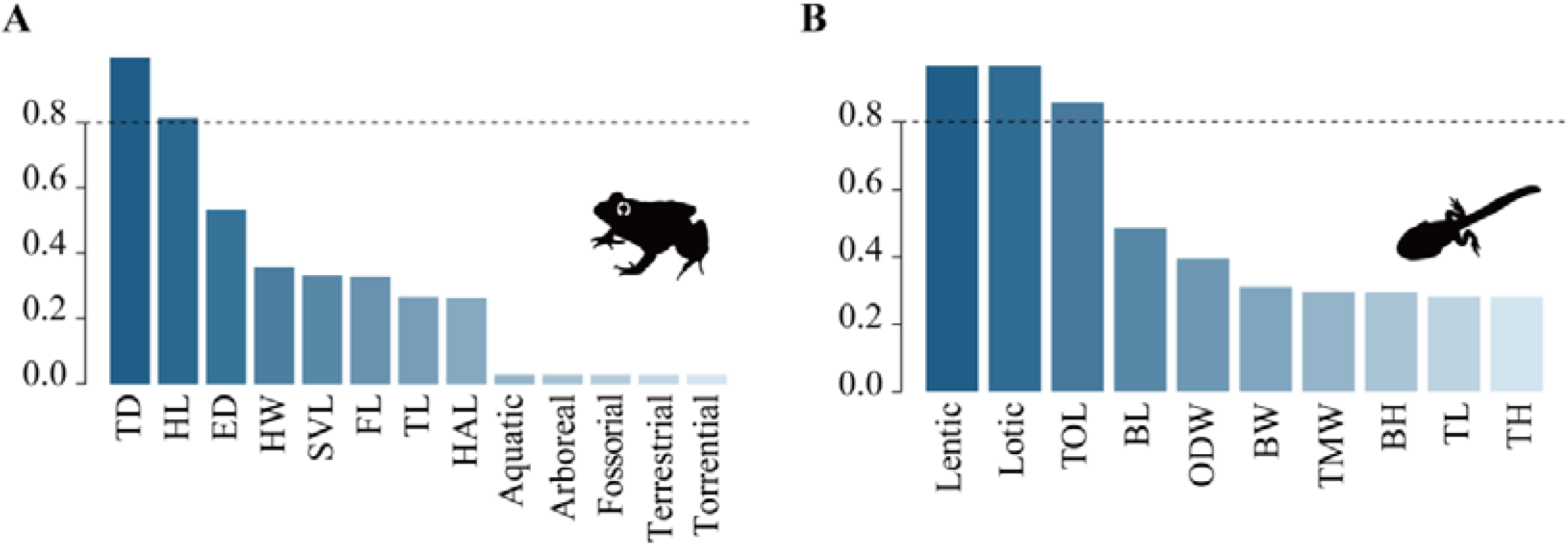
Trait importance (sum of Akaike Weights) of adult and tadpole traits based on model averaging under the OU evolutionary model using IUCN Red List. For abbreviations of traits, please refer to Table 1. The silhouette images are sourced from https://www.phylopic.org.

**Figure S5.**
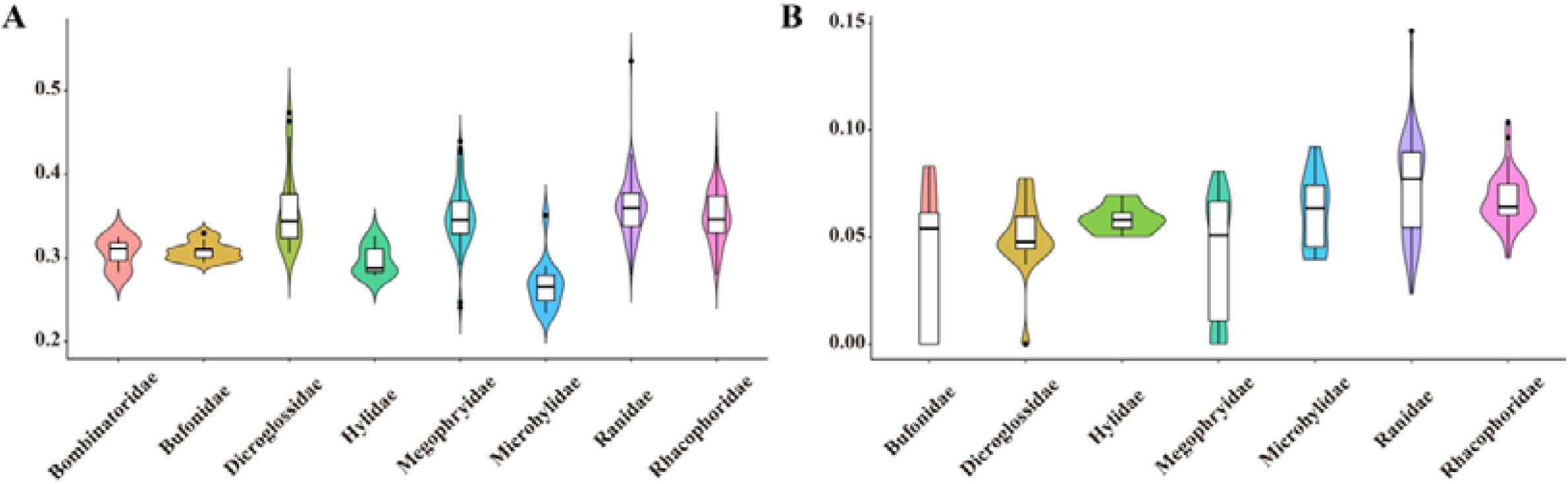
Violin plots showing the distribution of (A) head length and (B) tympanum diameter across anuran families in China. Species of the Bombinatoridae lack a tympanum; therefore, they are marked as 0 and are not shown in Figure B. Additionally, because other families also include species without a tympanum, the kernel density estimation produces boundary effects, leading to negative values in the plots. Thus, we truncated the density estimation at the data boundaries during visualization. Each violin plot represents the kernel density estimate of trait values, with the inner boxplot indicating median, interquartile range, and outliers. Traits are standardized relative to snout–vent length.

